# Integrative Network and Brain Expression Analysis reveals Mechanistic Modules in Ataxia

**DOI:** 10.1101/437582

**Authors:** Ilse Eidhof, Bart P van de Warrenburg, Annette Schenck

**Affiliations:** Department of Human Genetics, Donders Institute for Brain, Cognition, and Behavior, Radboud university medical centre, Nijmegen, The Netherlands.; Department of Neurology, Donders Institute for Brain, Cognition, and Behavior, Radboud university medical centre, Nijmegen, The Netherlands.

## Abstract

**Background:** Genetic forms of ataxia are a heterogenous group of degenerative diseases of the cerebellum. Many causative genes have been identified, but a systematic investigation of these genes to understand ataxia pathophysiology has not been performed.

**Methods:** A manually curated catalogue of 71 genes involved in disorders with progressive ataxias as a major clinical feature was subjected to an integrated gene ontology (GO), protein network, and brain gene expression profiling analysis.

**Results:** We found that ataxia genes operate in networks with significantly enriched protein connectivity, demonstrating coherence on a global level, independent of inheritance mode. Moreover, elevated expression specifically in the cerebellum predisposes to ataxia. Genes expressed in this pattern are significantly overrepresented among ataxia genes and are enriched for ion homeostasis/synaptic functions. The majority of ataxia genes, however, does not show elevated cerebellar expression that could account for region-specific degeneration. For these, we identified defective cellular stress responses as a major common biological theme, suggesting that the defense pathways against stress are more critical to maintain cerebellar integrity than integrity of other brain regions. Approximately half of the ataxia genes, mostly part of the stress module, show higher expression at embryonic stages, which argues for a developmental predisposition.

**Conclusion:** Genetic defects in ataxia predominantly affect neuronal homeostasis, to which the cerebellum appears to be excessively susceptible. Based on the identified modules, it is conceivable to propose common therapeutic interventions that target deregulated calcium and ROS levels, or mechanisms that can decrease the harmful downstream effects of these deleterious insults.

## Introduction

Genetic cerebellar ataxias are a group of disabling disorders that share progressive incoordination of movement due to dysfunction and degeneration of the cerebellum as their main hallmark^1^. The advent of next-generation sequencing technologies greatly advanced the identification of genes involved in ataxia^2^. However, despite the increasing number of genes identified, treatment attempts are still limited to relieving symptoms and do not target the underlying biological mechanisms. Development of effective therapies is hampered by an enormous genetic heterogeneity, the rarity of some the subtypes, and the limited knowledge of biological processes in which ataxia genes exert their function. Identification of shared biological modules between ataxia genes would provide a basis for therapeutic strategies that could be applied to larger cohorts of ataxia patients, in spite of their heterogeneous genotypic background.

In recent years, efforts have been made to identify common denominators of genetic ataxias. A number of ataxia genes were found to share interaction partners at protein level and to be involved in processes such as RNA splicing, regulation of transcription, and cell cycle^2–5^. When impaired in animal models, these processes lead to neurodegeneration, suggesting that these shared biological pathways maintain the integrity of the cerebellum and its connections^3^. Nevertheless, previous studies focused primarily on protein networks among specific ataxia genes and subtypes, and they did not systematically probe the influence of gene expression on cerebellar pathology^2–5^.

In this study, we systematically analyze the genes to date implicated in cerebellar ataxia, their functional biological pathways, and their expression in the developing human brain. Our integrative study identifies common denominators that underlie progressive cerebellar degeneration and ataxia, including a cerebellum-specific mechanism affected in a subgroup of ataxia disorders that may account for region-specific degeneration and defective stress defense pathways as underlying mechanism to the large majority of ataxia disorders to which the cerebellum is in particular sensitive.

## Materials and methods

### Cerebellar Ataxia gene selection and classification

The Human Phenotype Ontology database^6^ (download 11/2015) was used to search for genes associated with ataxia. A list of 347 genes was obtained and manually curated using PubMed and OMIM. Genes associated with progressive cerebellar ataxia as prominent clinical manifestation, either in isolation or as part of a more complex phenotype, were included. Primary metabolic disorders, genes associated with cerebellar hypoplasia, and genes inconsistently associated with cerebellar ataxia were excluded.

### Gene Ontology analysis

The webtool G-profiler^7^ (rev 1536, build 02/2016) was used to perform Gene-Ontology (GO) analysis of 4 categories of ataxia genes (all genes, dominant genes, recessive genes, polyQ genes; the latter refers to the group of dominant ataxias genes that, when mutated, carry a coding CAG repeat expansion that leads to polyglutamine expansion in the protein). For this study, we only considered GO terms (Biological Process, Molecular Function and Cellular Component) that were significantly enriched after correction for multiple testing (Bonferroni test, p<0.05). GO analysis for the developmental transcriptome data of BrainSpan was performed using the filtered gene-list (average RPKM > 0.05 over all developmental stages) as background.

### Enrichment analysis

Enrichment scores for GO-terms were calculated as followed: (*a*/*b*)/[(*c* − *a*)/(*d* − *b*)], where *a* is the number of genes in the ataxia category associated with that GO-term, *b* is the total number of genes in the ataxia category, *c* is the total number of genes in the genome associated with that GO-term or, for GO analysis of brainspan data, the total number of genes associated with that GO-term remaining after filtering out low expressed genes from BrainSpan data and *d* is the total number of annotated human genes present in Ensembl (20,313 genes) or, for GO analysis of brainspan data, the total number of genes remaining after filtering out low expressed genes from BrainSpan data (16,956 genes).

### Protein-protein interaction network

Protein-protein interactions (PPI) between CA genes were obtained from GeneMANIA, HPRD and BioGrid and included physical interactions, predicted interactions, shared protein domains, and pathways^8–10^. All interactions were combined and assembled in a reference network using Cytoscape^11^ (v3.1.1.) and duplicates were removed from the reference network.

### Physical interaction enrichment (PIE) score

We used the PIE algorithm to account for biases in the number of reported protein interactions for disease-associated genes in the generated reference PPI network^12^. PIE scores and associated p-values were calculated against 10,000 random protein groups obtained by number-matched sub-samplings selected from the reference PPI network for all four ataxia gene categories, as previously described^13^ ^14^.

### BrainSpan developmental transcriptome analysis

The publically available developmental transcriptome RNA sequencing (RNA seq) data from the Human BrainSpan atlas was used for ataxia gene expression analysis. BrainSpan provides RNA seq count data represented as reads per kilobase per million mapped reads (RPKM) of 11 targeted neocortical human brain regions and 5 targeted non-neocortical brain regions. Details on samples, sequencing protocols and RNA expression analysis can be found at the brainspan website (http://www.brainspan.org). First, low expressed genes, with an average expression <0.05 RPKM over brain developmental stages and regions, were filtered out. Data was then binned into nine stages, spanning important developmental milestones of the prenatal and postnatal human brain. EdgeR^15^ (version 3.16.5) and Limma^16^ (version 3.30.7), provided by the online service Bioconductor, were used to identify the differentially expressed genes between the 16 brain regions separately over the developmental stages. Genes were considered to be differentially expressed between two brain regions if the adjusted P-value passed the <0.05 threshold. Graphpad Prism 5.0 contigency tables and Chi-square with Yates correction (two-tailed) tests were used to calculate whether ataxia gene expression was significantly enriched in the cerebellum compared to the rest of the brain. GENE-E (version 3.0.215, www.broadinstitute.org) was utilized to hierarchically cluster average RPKM values of ataxia genes in the cerebellum over the nine defined developmental stages.

## Results

### A systematic catalogue of genes associated with progressive Cerebellar Ataxia

We generated a manually curated systematic catalogue of 103 disorders consistently associated with progressive cerebellar ataxia, corresponding to 71 annotated protein-coding genes (**Table S1**). Of the 71 genes that met our criteria, mutations in 42 of the genes follow recessive inheritance and mutations in 31 of the genes follow dominant inheritance. Two genes, *SPTBN2* and *AFG3L2*, have been described in both dominant and recessive ataxia. Generally, mutations associated with recessive ataxia are loss of function mutations, whereas dominant ataxias can be caused by a combination of gain and/or loss of function mutations. Several genetic ataxias are caused by unstable repeat expansions that can occur in noncoding and coding regions of the genome. Eight of these (*ATXN1*, *ATXN2*, *ATXN3*, *ATXN7*, *ATXN8*, *CACNA1A*, *TBP* and *ATN1*) contain/represent translated trinucleotide repeats that encode for polyglutamine (polyQ) residues and follow dominant inheritance.

### Genes involved in ataxia function in common biological processes

Differences in the type of mutations and phenotypes between recessive and dominant ataxias suggest that in spite of shared clinical features, different biological mechanisms might underlie these disorders^2 5^. To examine whether recessive, dominant and polyQ ataxia genes are associated with distinct biological functions, GO analysis was performed on the three gene categories. Recessive ataxia genes were significantly enriched for DNA metabolic processes (p=0.017), DNA-dependent DNA replication (p=0.038), DNA repair (p=4.2e^−4^), cellular response to stress (p=0.002), and mitochondrion (p=0.025) (**Fig 1A**). Dominant ataxia genes were significantly enriched for nuclear matrix (p=0.023), somatodendritic compartment (p=0.008) and dendrite (p=0.009) (**Fig 1B**). Ataxia genes encoding PolyQ expansions were significantly enriched for regulation of cellular biosynthetic process (p=0.047), nuclear periphery (p=8.57e^−5^), nuclear matrix (p=3.81e^−5^) and nuclear inclusion body (p=0.012) (**Fig 1C**). Despite these differences, the shared clinical hallmarks of genetic ataxias suggest that ataxia genes/proteins might affect common biological pathways or processes. To identify such common biological themes, we also analyzed the combined ataxia gene catalogue. We found a significant enrichment for most of the GO terms revealed by the separate analyses of genes underlying ataxia subtypes, including cellular response to stress (p=0.003) and DNA repair (p=0.001) (recessive), dendrite (p=0.003) (dominant), and nuclear inclusion body (p=0.029) (PolyQ) (**Fig 1D**). Interestingly, analysis of the contribution of recessive and dominantly inherited ataxia genes to the identified GO terms revealed a shared contribution to nearly all processes (**Fig 1D**), supporting an overlapping molecular pathology underlying both dominant and recessive ataxias. Moreover, several GO terms, such as those linked to calcium ion transmembrane transport (p=0.020), neuron projection (p=0.004), and adult walking behavior (p=0.014), were only highlighted if the analysis was applied to the complete catalogue of ataxia genes and might be more representative of the shared hallmarks between recessive and dominant progressive cerebellar ataxias.

**Figure 1.**
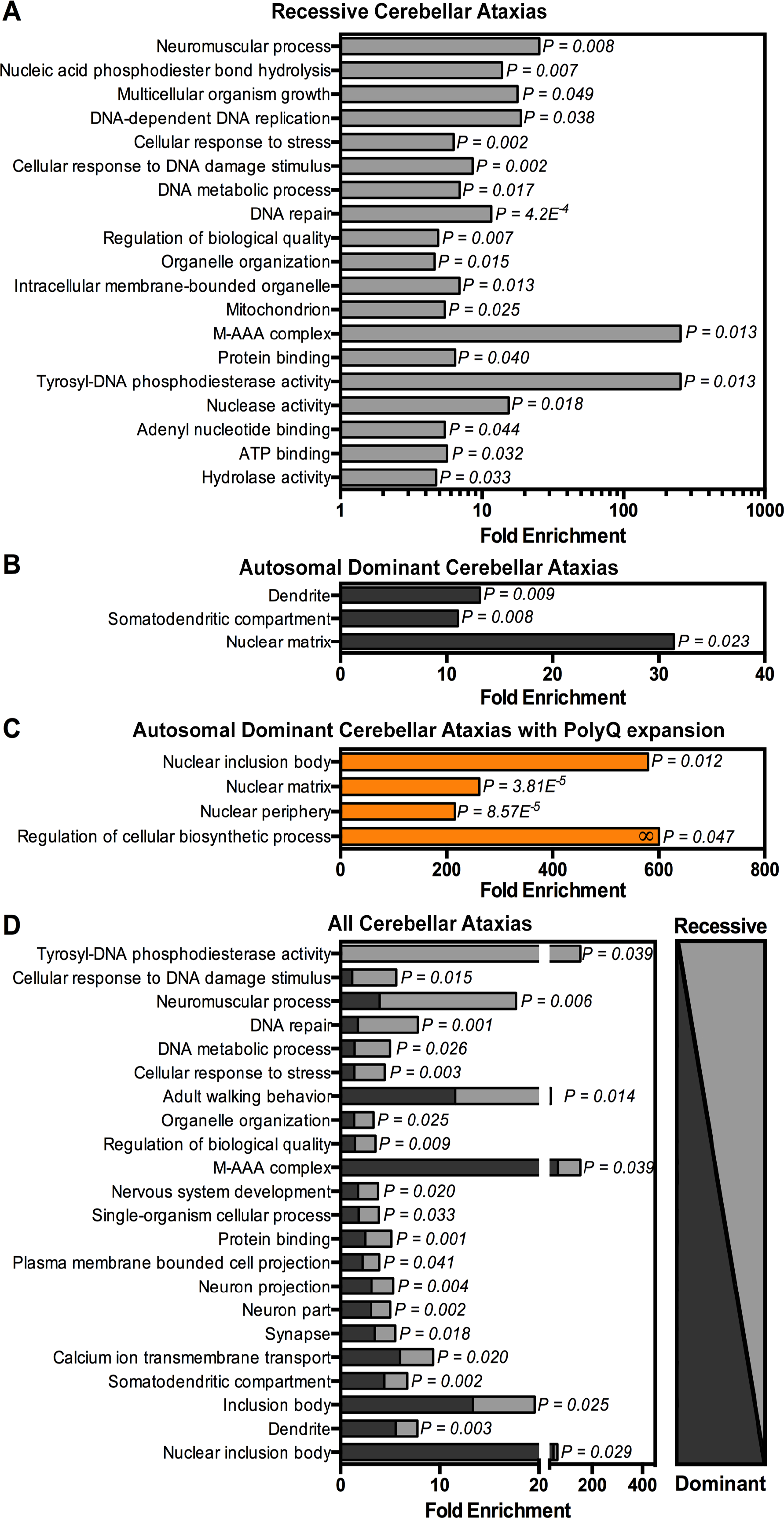
Cerebellar Ataxia genes function in common biological processes. GO-terms significantly enriched among (A) all ataxia genes, (B) Recessive ataxia genes. (C) Dominant ataxia genes, (D) Dominant ataxia genes with PolyQ expansion. (All GO-terms passed Bonferoni correction for multiple testing, p<0.05)

### Genes involved in ataxia show high connectivity on the protein level

Physical and functional interactions of proteins provide the basis of biological pathways and are crucial to understand cellular function. We evaluated whether our catalogue of ataxia proteins shows significant molecular connectivity. For this, we collected PPI data from three large databases and combined these into a reference network (**Fig 2A**). We found that 46 of the proteins physically interacted with other proteins present in the ataxia catalogue. Of these 46 proteins, 17 proteins were connected in small modules (pairs, tri- and hexamers), whereas 29 proteins formed a single major network with 30 interactions (**Fig 2A**). Interestingly, within these modules and networks, proteins associated with recessive, dominant and polyQ progressive ataxia are jointly represented, demonstrating a biological overlap regardless of inheritance type or mutational mechanism (**Fig 2A**

To assess the significance of the identified connectivity, PIE scores and associated p-values were calculated for all, and separately for recessive, dominant and polyQ ataxia proteins. This analysis revealed that ataxia proteins as a whole group interact 2-fold more than randomly expected (P<0.001) (**Fig 2B**). Recessive ataxia proteins interacted 2.1-fold more (p<0.001), dominant ataxia proteins interacted 2.8-fold more (p<0.001), and polyQ-associated ataxia proteins interacted 3-fold more (p<0.05) (**Fig 2B**). Thus, ataxia proteins are significantly interconnected.

**Figure 2.**
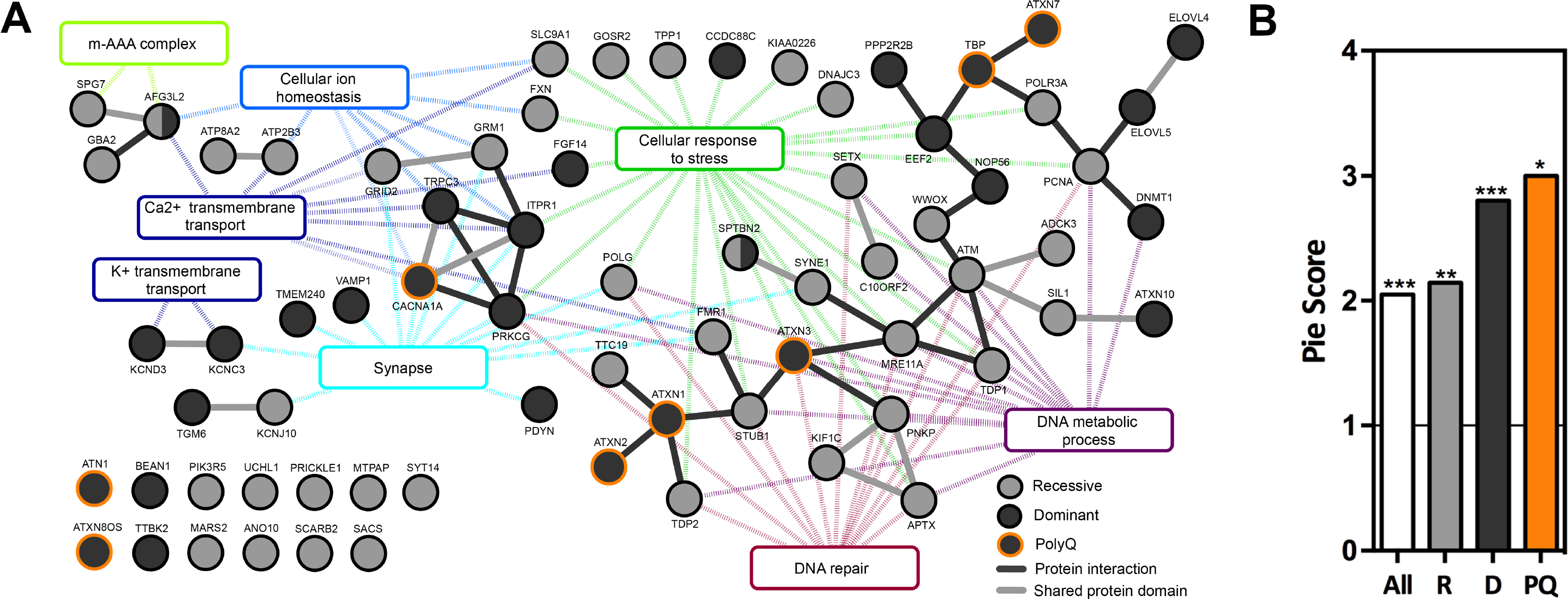
Cerebellar Ataxia proteins show high connectivity on protein level and function in common processes. (A) Interaction network of ataxia proteins. (black solid lines: direct protein interaction, grey solid lines: proteins with similar domains, dotted lines: interaction of protein with GO-term) (B) PIE Score of All, Recessive (R), Dominant (D) and PolyQ (PQ) ataxia proteins. (*** P<0.001, ** P<0.01, * P<0.05, based on 10,000 random repetitions)

Since interactions between proteins underlie biological processes and pathways, we continued examining whether these represent specific biological processes. Individual small protein modules were significantly enriched for processes such as calcium ion homeostasis (p=0.001) and calcium ion transmembrane transport (p=0.024), ATP-dependent peptidase activity, m-AAA complex (p=4.27e^−6^), unfolded protein binding (p=0.024), potassium ion transport (p=0.024), and potassium channel activity (p=0.007) (**Fig 2A**, **left side**). The large module within the ataxia interactome was significantly enriched for DNA repair (p=3.23e^−5^), cellular response to stress (p=0.020), and nuclear inclusion body (p=0.001) (**Fig 2A**, **right side**). Together, these results illustrate that ataxia proteins function in common biological networks and processes.

### A subset of ataxia genes shows high relative expression specifically in the cerebellum

The basis for the preferential regional vulnerability of neurons in the cerebellum in genetic ataxias is mostly unknown. Temporal and spatial patterns of ataxia gene expression in the brain may significantly contribute to specific or preferential cerebellar degeneration when disturbed. To address this, we turned to the publicly available BrainSpan Transcriptional Atlas of the Developing Human Brain, as resource. After exclusion of low expressed genes, a transcription matrix of 16,956 genes representing the 16 brain regions was left that was binned into nine different developmental stages (**Fig 3A**). We then calculated for each of the 16,956 genes per developmental period whether it was differentially expressed in the cerebellum, compared pairwise to any of the other 15 brain regions. From the resulting matrix, the ataxia genes were extracted and the percentage of ataxia genes differentially expressed (adj. p<0.05) in the cerebellum compared to one or more other brain regions was calculated and visualized in a heatmap (**Fig 3B**). Of note, the expression levels of three ataxia genes, *TGM6*, *PIK3R5* and *MTPAP*, did not pass the threshold of very low expressed genes and were therefore excluded from further analyses. The expression of most ataxia genes in the cerebellum was not different from any of the other 15 brain regions. However, the vast majority of those ataxia genes that did show differential expression to other brain regions, did so to all 15 of them. Their high relative expression level was specific to the cerebellum, suggesting their distinct requirement in this region of the brain. Interestingly, this pattern showed a sharp onset at birth, suggesting that these genes serve cerebellar function specifically at postnatal stages (**Fig 3B**).

**Figure 3.**
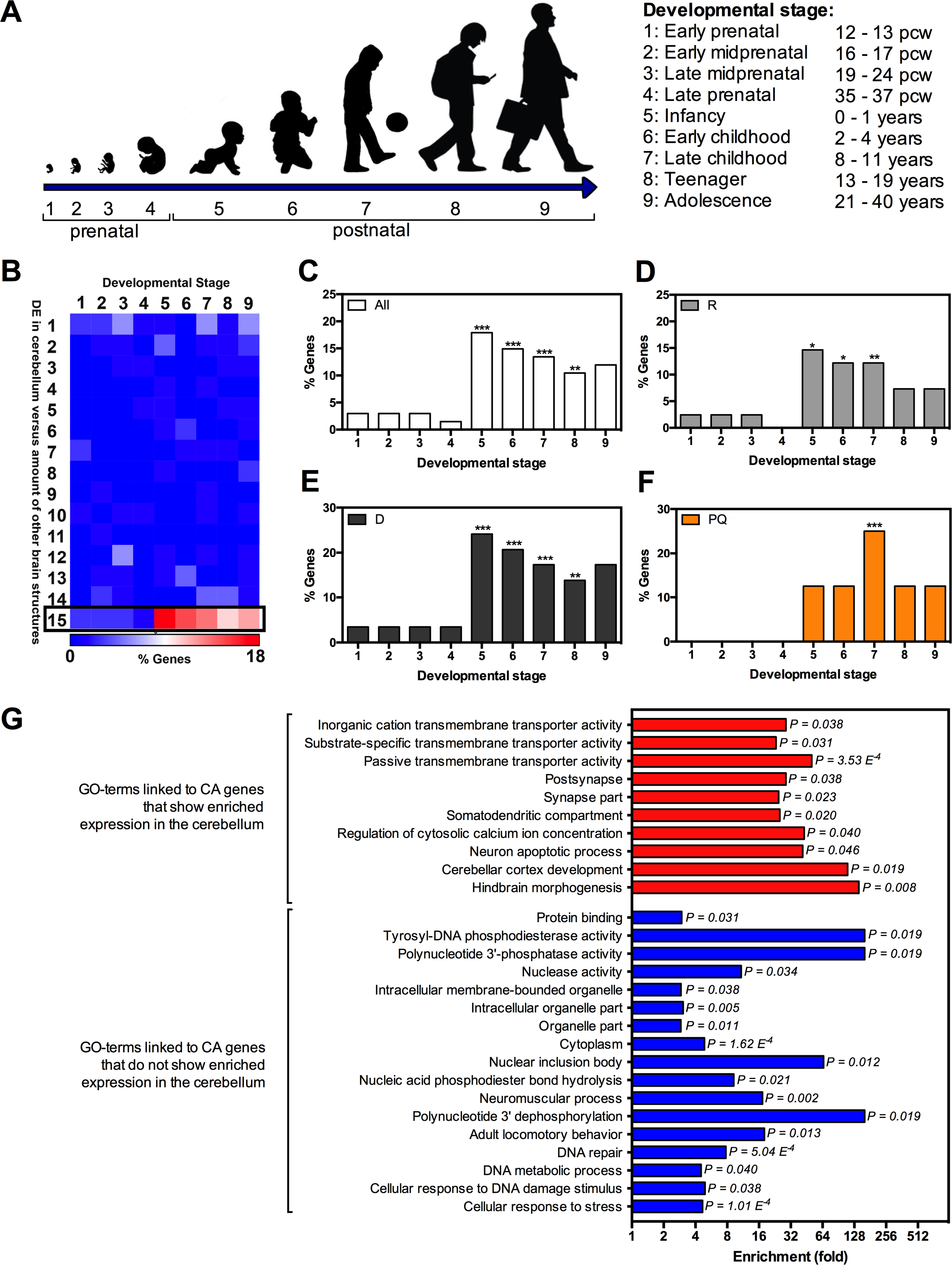
Cerebellar Ataxia gene expression is enriched in the postnatal cerebellum. (A) 9 developmental stages were used for analysis of developmental human BrainSpan expression data. (B) Heatmap displaying % of ataxia genes differentially expressed in the cerebellum compared to indicated amount of other non-cerebellar brain regions for developmental stage 1-9. (C-F) % genes that show significant enriched expression in the cerebellum compared to 15 other brain regions for described developmental stages ((C) All ataxia genes, (D) Recessive ataxia genes, (E) Dominant ataxia genes, (F) Dominant ataxia genes with PolyQ expansion, * P<0.05, ** P<0.01, ***P<0.001). (G) Significantly enriched GO-terms for ataxia genes elevated in the cerebellum (red) and ataxia genes not elevated in the cerebellum (blue).

### Elevated cerebellar-specific expression predisposes to ataxia

We continued determining whether elevated relative expression in the cerebellum predisposes to ataxia. To address this, we calculated whether the amount of ataxia genes that were specifically elevated in the cerebellum for each developmental stage was higher than randomly expected, considering the genome-wide frequency of cerebellar elevated genes. Ataxia genes were more frequently elevated in the cerebellum during the postnatal stages 5-9 (infancy: p<0.001, early childhood: p<0.001, late childhood: p<0.001 and teenager: p<0.01) than randomly expected (**Fig 3C**). The following genes were specifically elevated in the cerebellum during one or more of the analyzed developmental stages: *ADCK3*, *ATM*, *ATN1*, *CACNA1A*, *DNMT1*, *GRID2*, *GRM1*, *ITPR1*, *KCNC3*, *KCND3*, *SPTBN2*, *SYNE1* and *TRPC3* (**Table S1**). To further examine whether a specific mode of inheritance or mutational mechanism was underlying this group of genes, the analysis was repeated for the recessive, dominant and polyQ ataxia gene categories. Also here, the expression of genes involved in either of these three categories was enriched in the cerebellum during postnatal development stages (**Fig 3D-F**), suggesting that all of them contribute to the finding of cerebellum-specific postnatal enrichment of ataxia gene expression.

Finally, we asked whether the genes that were not specifically elevated in the cerebellum during one of the nine developmental stages, were specifically elevated in one of the other 15 brain regions. We found that a small group of ataxia genes was specifically elevated in either the thalamus (*GBA2*, *UCHL1*, *PRICKLE1* and *ANO10*) or striatum (*CCDC88C* and *PDYN*) during certain developmental stages compared to rest of brain. Not a single ataxia gene was significantly elevated in one of the remaining 13 specific brain regions.

In summary, systematic analyses of brain expression across brain regions revealed that the expression of a subgroup of ataxia genes is specifically enriched in the postnatal cerebellum, likely driving the pathological features of these disorders.

### Cerebellar-specific expression patterns separate ataxia genes in distinct biological modules

We next asked which functional biological modules are underlying the identified group of genes with cerebellar-specific expression. GO term analysis was performed on the lists of ataxia genes that showed enriched cerebellar expression during one or more of the developmental stages and ataxia genes that did not (**Table S1**). This revealed that ataxia genes that were specifically elevated in the cerebellum during one of the developmental stages were significantly enriched for neuronal and ion related processes such as: neuron apoptotic process (p=0.046), hindbrain morphogenesis (p=0.008), regulation of cytosolic calcium ion concentration (p=0.040), synapse part (p=0.023), and passive transmembrane transporter activity (p=3.53e^−4^) (**Fig 3G**). Ataxia genes that were not specifically elevated in the cerebellum during one of the developmental stages were significantly enriched for processes such as: DNA repair (p=0.004), cellular response to stress (p=0.023) and nuclear inclusion body (p=0.035) (**Fig 3G**).

### Temporal gene expression patterns in the cerebellum cluster genes involved in progressive ataxia into two groups

Finally, we also explored developmental expression profiles of ataxia genes in the cerebellum over time, to examine whether this provides further clues about dependence of the cerebellum on certain biological processes during specific stages of development (**Fig 4A, B**). We hierarchically clustered the temporal expression patterns of ataxia genes in the cerebellum over the nine developmental stages, which unbiasedly separated the genes in two distinct clusters (**Fig 4B**). Genes present in cluster 1 showed significant higher expression during prenatal stages compared to genes in cluster 2, and genes present in cluster 2 showed significant higher expression during postnatal stages compared to genes present in cluster 1 (**Fig 4A**). Recessive, dominant and polyQ associated ataxia genes contributed randomly to the two clusters (data not shown). However, genes expressed higher during prenatal stages (cluster 1) were enriched for DNA repair (p=0.031), whereas genes with higher postnatal expression (cluster 2) were enriched for processes related to the cellular component synapse (p=0.019), calcium ion transmembrane transport (p=0.012), and metal ion homeostasis (p=0.042) (**Fig 4C**). Ataxia genes can thus be distinguished based on their temporal expression profiles in the cerebellum, which links to specific biological processes.

**Figure 4.**
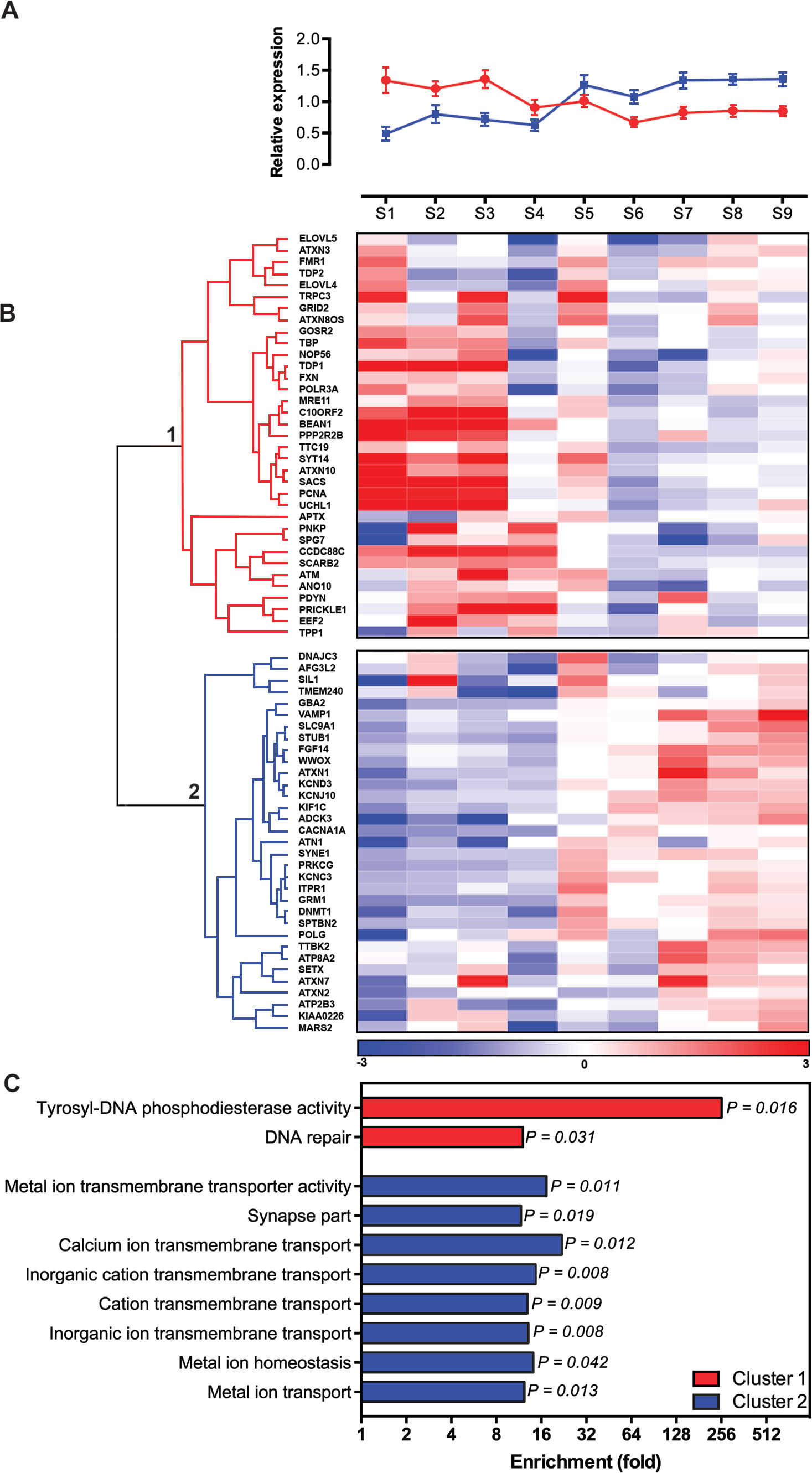
Ataxia genes can be separated in two distinct clusters based on their temporal expression levels in the cerebellum. (A) Average RPKM value for ataxia genes present in cluster 1 and cluster 2 of Fig 4B, for the indicated developmental stages. (pink: cluster 1, blue: cluster 2, error bars represent standard deviation) (B) Hierarchical clustering of ataxia gene expression levels during cerebellar development using Spearman correlation. Data was obtained from BrainSpan and mean RPKM values were calculated for the indicated developmental stages. Heatmap color-codes are based on median RPKM value per row (developmental stage), divided by the row standard deviation (blue: low expression in cerebellum compared to median, red: high expression in cerebellum compared to median). (C) Significantly enriched GO-terms for Cluster 1 and Cluster 2 from Fig 5B (All GO-terms passed Bonferoni correction for multiple testing, p<0.05).

## Discussion

We have here systematically mapped shared molecular pathways, processes and expression characteristics among ataxia causing genes, to increase our understanding of the biology of genetic ataxias and to identify mechanistic hubs that can serve as targets for therapeutic interventions. In comparison to previous studies^2–5^; we generated a manually curated catalogue of genes involved in genetic ataxia and performed analyses across dominant and recessive forms of the disorder.

Furthermore, we applied a strategy that integrated ataxia gene expression in the developing human brain, gene ontology, and protein interaction network analysis, to get a comprehensive understanding of the vulnerability of the cerebellum and the molecular modules and processes affected in genetic ataxias.

Data in this study highlight the different biological processes that are implicated in recessive, dominant and polyQ ataxias, with recessive ataxias linked to cellular response to stress and DNA repair related processes; dominant ataxias to dendrite related processes; and polyQ ataxias to nuclear inclusion body. The deleterious effect of the type of mutation on the protein, the expression timing profile, specificity and levels of the affected protein in the cerebellum, and the sensitivity of the cerebellum to disruption of these processes might explain these different findings. Despite of this, we found a shared contribution of recessive and dominant ataxia genes to nearly all biological processes, while processes such as calcium ion transmembrane transport were only enriched when applying GO analysis to the complete ataxia gene panel.

We found that ataxia genes operate in networks with significantly enriched protein connectivity, demonstrating global coherence independent of inheritance mode or mutational mechanism. Notably, polyQ proteins interacted directly with other non-polyQ dominant and recessive ataxia proteins. This indicates that, in addition to common toxic gain-of-function mechanisms^17^ such as the formation of nuclear inclusion bodies, the disruption of the biological processes that these genes operate in likely contributes to the disease pathogenesis. This notion derived from our systematic analysis is supported by gene-focused studies, e.g. of mice models of SCA1, where ATXN1 loss-of-function phenotypes were very similar to ATXN1 gain-of-function phenotypes, and of SCA17 models demonstrating that impaired transcriptional activity of polyQ-expanded TBP contributes to disease pathogenesis^18–20^. Together, our findings strongly support an overlapping molecular pathology between recessive and dominant ataxia subtypes.

The identified protein modules represent different biological processes and ataxia proteins can broadly be separated in two themes: a large stress module and smaller ion homeostasis/synapse modules. The common end-point of these modules is progressive degeneration of the cerebellum, and analysis of ataxia gene expression in the developing human brain suggested that these two modules might contribute differently to cerebellar vulnerability, depending on their specific and temporal cerebellar expression pattern (**Fig 5**).

**Figure 5.**
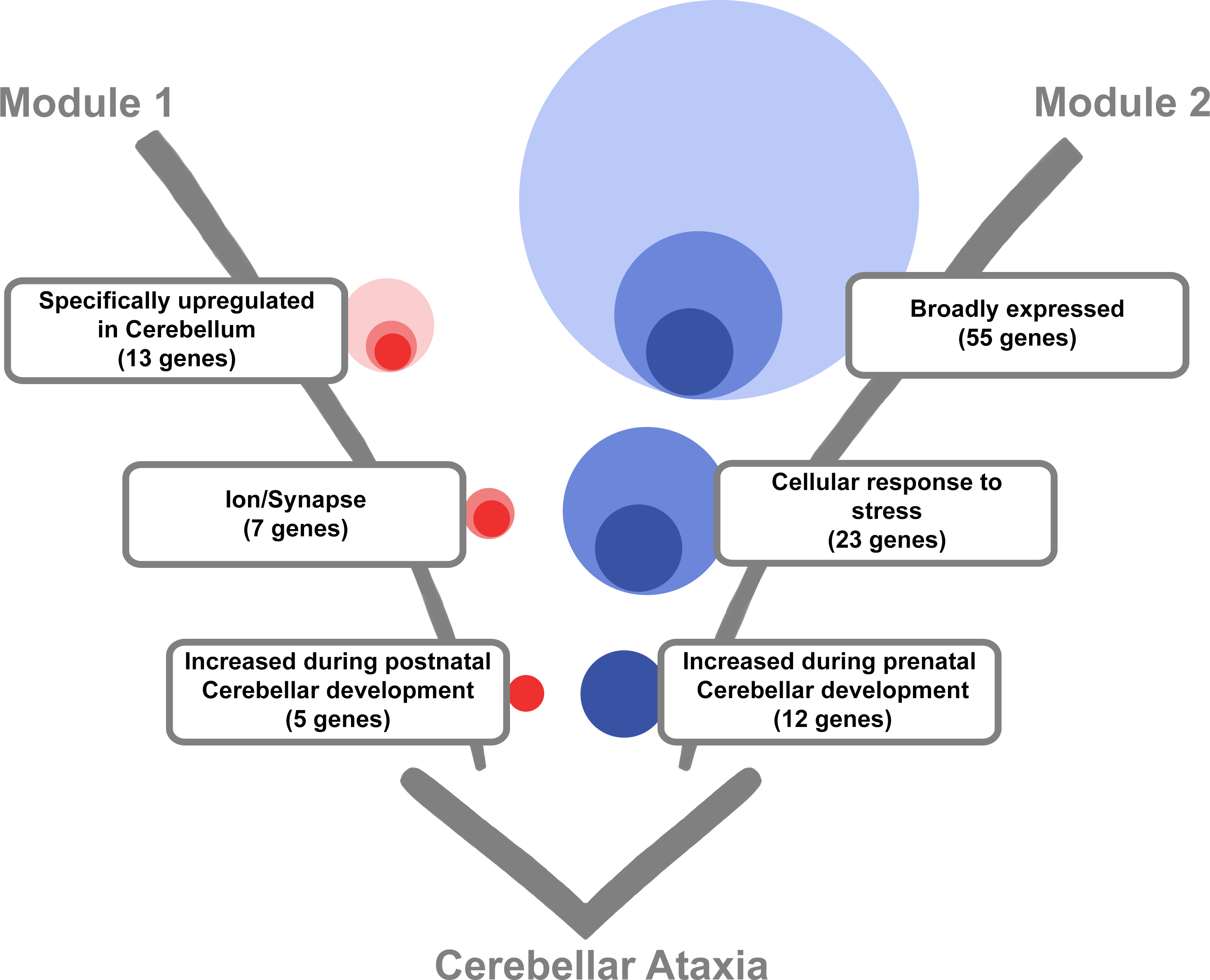
Ataxia genes can broadly be divided in two themes that affect neuronal homeostasis and when disrupted predispose to progressive cerebellar ataxia. Module 1: 13 Genes are specifically elevated in the cerebellum during one or more developmental stages. Of these 13 genes, 7 are linked to Ion/Synapse function and 5 out of these 7 genes showed increased expression during postnatal cerebellar development compared to prenatal cerebellar development. Module 2: 55 genes do not show increased expression in the cerebellum. Of these 55 genes, 23 genes are linked to cellular response to stress and 12 out of these 23 genes show increased expression during prenatal cerebellar development compared to postnatal cerebellar development (data displayed in Fig 2-4).

Brain expression analysis demonstrated that elevated gene expression specifically in the postnatal cerebellum predisposes to ataxia. Genes with this pattern of expression are significantly overrepresented among ataxia genes. The normally high gene activity levels may explain the specific vulnerability of the cerebellum to deleterious mutations in these genes. Interestingly, the corresponding genes encode ion channels, calcium receptors, calcium-activated proteins enriched for transmembrane transporter activity, and regulators of cytosolic calcium ion concentration. Most of them are part of the smaller ion homeostasis/synapse module we identified (**Fig 3**). The cerebellum contains a unique neuronal cell type, the Purkinje cell (PC), that is central to ataxia pathobiology. PCs might be particularly vulnerable for alterations in ion homeostasis due to their extensive dendritic arbor that exhibit intense and highly regulated firing properties^21–23^. The striking complexity of PC firing regulates sensorimotor integration and is highly dependent on calcium channels and calcium-activated potassium channels^21^. In ataxia models, PCs show reduced firing rate and loose intracellular calcium buffers, even before the onset of clinical symptoms and PC degeneration^17^ ^24^. Reduced PC excitation enhances the excitability of efferent deep cerebellar nuclei neurons and this is sufficient to cause cerebellar ataxia even in the absence of PC degeneration^25^. This shows that calcium homeostasis in PCs is crucial for proper sensorimotor integration and disruption of these processes likely affects PC firing properties, eventually leading to cerebellar degeneration and the onset of ataxia.

Calcium is also an important messenger in intracellular signaling pathways, proteostasis mechanisms at ER and Golgi membranes, and mitochondrial activity^26^ ^27^. The metabolic activity of PCs is high^28^. Deficits in neuronal energy production and intracellular organelle systems that influence ion fluxes may thus be further factors that account for increased vulnerability of PCs. The energy provided for metabolism is provided by mitochondrial oxidative phosphorylation in the form of adenosine triphosphate. Increased calcium uptake by mitochondria leads to extensive oxidative phosphorylation and overproduction of Reactive Oxygen Species (ROS)^26^ ^27^. ROS overproduction in turn can lead to detrimental oxidative modifications of lipids, proteins and nucleic acids^29^ ^30^. Interestingly, the large identified protein module is enriched for the cellular stress response, and proteins present in this module are among others involved in mitochondrial maintenance, DNA repair, unfolded protein response, and regulation of apoptotic and autophagic processes. We found that overall, these genes were not more elevated in the cerebellum during any of the analyzed developmental stages. Thus, elevated cerebellar gene expression cannot account for preferential cerebellar degeneration. This suggests that, compared to other neuronal cell types, the PCs might be more vulnerable to insults related to a distorted cellular stress response. Interestingly, approximately half of the cellular stress response genes show higher expression during embryonic stages, arguing for a developmental predisposition in these cases.

Ataxia genes with a function in DNA repair were abundantly represented in the stress module (**Fig 3**). DNA breaks arising from oxidative damage are a major threat for the genome stability of mature post-mitotic neurons. They are usually repaired by base excision repair and single strand break repair (SSBR)^31^. Interestingly, genetic ataxias such as SCAN1 and AOA1, are associated with DNA SSBR deficiencies, and animal models of these show increased sensitivity to ROS^28^ ^31^. The genes mutated in these disorders, *TDP1* and *APTX*, do not show enriched cerebellar expression during any of the nine analyzed developmental stages, which may indicate that SSBR is a key homeostatic process required in PCs for reasons unrelated to expression, such as their high metabolic activity, oxidative load, and intrinsic firing properties. There is also a distinct subset of genes implicated in recessive ataxias that are involved in Double Strand Break (DSB) repair. Double stranded DNA breaks commonly occur during rapid proliferation of CNS development and are less likely to occur in the matured nervous system^32^. The cerebellum might be in particular vulnerable to disruption of DSB repair due to its development up until the first postnatal years, that will lead to the formation of the most abundant cell type in the central nervous system: the cerebellar granule neurons^28^. This postnatal period of rapid and massive cell proliferation may generate replication stress-associated DNA damage that might affect the cerebellar granule neurons, and indirectly other cerebellar cell types such as the PCs to which they signal^28^. Interestingly, ATM, a kinase that is involved in detection of DSBs and is mutated in ataxia telangiectasia (AT), shows enriched expression in the cerebellum exactly during this postnatal developmental period. Since ATM is involved in DSB-induced apoptotic signaling, dysfunctional neurons may fail to be efficiently eliminated in the early AT cerebellum, and degenerate only later on^31^ ^33^ ^34^. This is in agreement with the early neurological problems and loss of the cerebellar granule neurons, the molecular cell layer, and PCs in AT^31^ ^35^ ^36^. Nonetheless, since ROS-induced DNA damage can also include DSBs and antioxidant treatment can promote the survival of cultured ATM-deficient PCs, DSB repair might also be required for cerebellar maintenance^31^ ^37^. Therefore, DSB repair is likely a key homeostatic process that maintains PC integrity.

In conclusion, while a number of molecular processes are involved in progressive cerebellar ataxia pathology, these intersect and form a common end-point: disrupted neuronal homeostasis, to which the cerebellum is either exclusively susceptible, or more than other brain regions. More experimental data are required to understand the dependence of the cerebellum on different aspects of neuronal homeostasis, such as calcium signaling, ROS, and DNA repair, particularly in absence of region-specific expression levels. However, based on the here identified biological themes, it seems conceivable to propose therapeutic interventions that target deregulated calcium and ROS levels, or mechanisms that can decrease the harmful downstream effects of these deleterious insults.

## Acknowledgements

We thank P. Cizek for the PIE score script and C. Gilissen for advice on the expression analysis. This research was supported by the E-RARE-3 Joint Transnational Call grant “Preparing therapies for autosomal recessive ataxias” (PREPARE; ZonMW 9003037604 to B.v.d.W. and A.S.) and by a Radboud university medical centre junior researcher grant.

## Supporting Information

**Table S1:**Processed expression data used for analysis of ataxia gene expression in the developing human brain.

